# Evaluating non-lethal tissue suitability for telomere length measurement in the Japanese eel

**DOI:** 10.64898/2026.05.09.723945

**Authors:** Yuta Moriguchi, Satoko S. Kimura, Manabu Kume, Junichi Takagi, Yoshinobu Uno, Junko Kanoh, Hiromichi Mitamura

**Author notes:** (Yoshinobu Uno): Graduate School of Technology, Industrial and Social Sciences, Tokushima University, Tokushima, Japan.

## Abstract

Telomere length (TL) is increasingly used in ecology as a biomarker of individual quality and environmental stress, yet research on non-model species with complex life histories remains limited. Because TL varies among tissues and across ages in a species-specific manner, identifying non-lethal tissues that reliably reflect whole-organism telomere dynamics is essential for longitudinal telomere studies in the field. This study aimed to evaluate tissue-specific TL in Japanese eel (*Anguilla japonica*), an endangered catadromous fish. We first mapped the chromosomal distribution of telomeric sequences using fluorescent in situ hybridization (FISH), the first application of this method in this species. We then tested whether muscle and caudal fin, which can be sampled easily and non-lethally, can serve as suitable proxy tissues for TL measurements in wild individuals. Relative telomere length (RTL) was quantified by qPCR in blood, brain, caudal fin, gonads, heart, liver, and muscle. FISH analysis confirmed telomeric repeats at all chromosomal ends, with only weak interstitial signals on three chromosomal pairs unlikely to affect qPCR-based estimates. A generalized additive mixed model and Wilcoxon’s signed-rank tests revealed significant inter-tissue differences: RTL was shortest in the brain and muscle and longest in liver, blood and caudal fin. Muscle and caudal fin RTL were significantly correlated with RTL in many other tissues, supporting their use as proxy tissues for longitudinal TL monitoring, including responses to environmental variation. Both total length and age were tested as explanatory variables for RTL, and the model including total length showed a better fit than the age-based model. Non-linear relationships between RTL and total length observed in several tissues suggest physiological shifts associated with growth and sexual differentiation. Overall, these findings advance understanding of telomere dynamics in eels and establish muscle and caudal fin as suitable tissues for repeated, non-lethal TL assessment in ecological and conservation contexts.

## Introduction

Telomeres are nucleoprotein complexes composed of tandem repeats of specific nucleotide sequences and associated proteins located at the ends of eukaryotic chromosomes [1]. In most vertebrates, the repeat sequence is (TTAGGG)_n_ [2]. Telomere length (TL) shortens with DNA replication [3] and oxidative stress [4, 5]. Conversely, TL can be maintained or elongated by telomerase [6]. In recent years, TL and telomere dynamics have been associated with environmental stress, reproductive performance, and survival rate [7–12], suggesting their potential as indicators of individual quality [e.g., 7] and biomarkers in ecology and conservation biology [10, 13].

Selecting an appropriate tissue is critical when using TL as a biomarker [14]. In mammals, birds, and reptiles, TL is commonly estimated from blood DNA [15–18], owing to its relatively low invasiveness and ease of collection [18]. In humans [19, 20], dogs (*Canis lupus familiaris*) [21], pigs (*Sus scrofa domesticus*) [22], zebra finches (*Taeniopygia guttata*) [23], and the lizard (*Ctenophorus pictus*) [18], blood TL has been reported to positively correlate with those in other tissues, supporting blood as a useful proxy. In contrast, blood TL in the Egyptian fruit bat (*Rousettus aegyptiacus*) showed no significant correlations with other tissues [24]. Wing TL was significantly correlated with TL in multiple tissues and can be sampled repeatedly with low invasiveness, suggesting that it represents a more suitable proxy for TL in this species [24]. These results highlight that among-tissue variation in TL can be species-specific.

In teleosts, fins and muscle are frequently used for TL measurement due to accessibility [e.g., 25–27], yet inter-tissue correlations vary widely among species. For example, fin TL correlates with those in other tissues in wild brown trout (*Salmo trutta*) [28], and TL correlations have also been reported between specific tissue pairs in medaka (*Oryzias latipes*) [29], whereas correlations are absent in aquaculture Atlantic cod (*Gadus morhua*) [30]. These contrasts highlight the need to characterize tissue-specific TL patterns in diverse species, such as non-model fishes, inhabiting natural, variable environments [10, 24].

Here, we focus on Japanese eel (*Anguilla japonica*), a long-lived catadromous fish that spawns in the western Mariana region and grows in East Asian rivers during 5 to more than 10 years until silvering [31–33]. The abundance of Japanese eel has declined since the 1970s, and thus the species has been listed as Endangered on the IUCN Red List since 2014 [34, 35]. The degradation and loss of freshwater habitats are considered one of the major factors of its decline [36, 37]. Because physiological condition can directly affect individual survival and growth, assessing the condition of Japanese eels in freshwater habitats is crucial for their conservation. For field-based applications of TL studies in eels, muscle and fins are promising candidate tissues as they can be obtained non-lethally and relatively easily, and regenerate readily, potentially allowing repeated sampling of the same individuals for longitudinal analyses of telomere dynamics. To measure TL from such samples, qPCR is currently the most widely used method because it requires only a small amount of DNA and allows rapid quantification [38, 39]. However, if interstitial telomeric sequences (ITSs), telomeric-like repeats located at internal chromosomal regions, are abundant within chromosomes, qPCR-based estimates can be biased [40]. Therefore, it is important to confirm the presence and the abundance of ITSs before applying qPCR in a given species.

In this study, we aim to establish a foundation for telomere research in Japanese eel by evaluating suitable tissues for non-lethal TL assessment. To support the interpretation of qPCR-based estimates, we first mapped the chromosomal distribution of telomeric sequences in farmed individuals using FISH to assess the potential influence of ITSs. We then compared TL in non-lethal, easily accessible tissues (muscle and caudal fin) with that in internal tissues (blood, brain, gonads, heart, and liver) from wild individuals to evaluate the suitability of these tissues as practical proxies for individual TL. Finally, we examined cross-sectional variation in TL with body size and age to describe the patterns of telomere dynamics in this endangered species.

## Materials and methods

### 2-1. Fluorescence in situ hybridization (FISH) using farmed individuals

Cell culture was performed as described previously [41]. Heart and peritoneal membrane were excised from farmed Japanese eels (*N* = 6, Omori Tansui Co., Ltd., Miyazaki, Japan) after anesthesia with 0.1% diluted 2-phenoxyethanol (FUJIFILM Wako Pure Chemical Corporation, Osaka, Japan), and primary fibroblast cultures were established. Fresh tissues were washed in Dulbecco’s modified Eagle’s medium (Thermo Fisher Scientific-GIBCO, Carlsbad, CA, USA) containing 5% antibiotic–antimycotic solution (Thermo Fisher Scientific-GIBCO). The washed tissues were then finely minced with sterilized scissors and plated on a collagen I-coated culture dish (AGC Techno Glass, Shizuoka, Japan), and cultured in Dulbecco’s modified Eagle’s medium supplemented with 15% FBS, 1% antibiotic–antimycotic solution, 2 ng/mL EGF, 2 ng/mL bFGF, and 1% ITS-G (all from Thermo Fisher Scientific). Cultures were incubated at 26°C in a humidified atmosphere of 5% CO_2_. Primary fibroblasts were harvested, and subcultured; no more than 10 passages were used for subsequent procedures. After passaging, cultured cells were treated with colcemid (Nacalai Tesque, Inc., Kyoto, Japan) at a final concentration of 150 ng/mL and incubated for 1–2 h, subjected to hypotonic treatment in 0.075 M KCl for 20 min, and fixed in Carnoy’s fixative (methanol:acetic acid = 3:1). The cell suspensions were dropped onto clean glass slides, air-dried at room temperature, and the chromosome preparations were stored at −80°C until use.

FISH was performed as described previously [41, 42]. Chromosome slides were hardened at 65°C for 2 h, denatured in 70% formamide/2× SSC at 70 °C for 2 min, and dehydrated in cold 70% ethanol (4°C) and 100% ethanol at room temperature for 5 min each. DIG-labelled 42 bp-long oligonucleotide sequences, (TTAGGG)_7_ and (TAACCC)_7_, were diluted in a hybridization mixture composed of 50% formamide/2× SSC, 10% dextran sulfate, BSA (2 μg/μL), and water at a 1:2:1:1 ratio, applied onto denatured chromosome spreads, covered with Parafilm, and incubated overnight at 37°C. After hybridization, the slides were washed in 50% formamide/2× SSC at 37°C for 20 min, then in 2× SSC at room temperature for 15 min, followed by 1× SSC for 15 min, and rinsed in 4× SSC for 5 min. For detection, slides were incubated under Parafilm with rhodamine-conjugated anti-DIG Fab fragments (Roche Diagnostics, Basel, Switzerland). The slides were then washed sequentially for 10 min each in 4× SSC, 0.1% Nonidet P-40 in 4× SSC, and 4× SSC, rinsed in 2× SSC for 5 min, and mounted with Vectashield mounting medium containing DAPI (Vector Laboratories, Burlingame, CA, USA). The digital FISH images were captured using a DeltaVision microscope system (Cytiva, Tokyo, Japan) and processed by deconvolution using SoftWoRx software (version 7.0.0, Cytiva).

### 2-2. Field sampling for tissue comparison in TL of wild individuals

To assess tissue-specific TL under natural conditions, wild yellow Japanese eels (*N* = 50) were collected from the lower reaches of the Iroha River, Oita Prefecture, Japan (Fig 1) on November 7, 2023, using an electrofisher (LR-20B, SmithRoot, Vancouver, WA, USA). No releases of farmed eels into this river have been reported. The collected individuals were transported alive under refrigerated conditions to the laboratory on the following day. Two days after capture, specimens were dissected after anesthesia with 0.1% diluted 2-phenoxyethanol for tissue collection. Blood, brain, caudal fin, gonads, heart, liver, and muscle were sampled, and stored at −30°C. Whole blood was collected into microtubes containing sodium heparin (Nacalai Tesque) as an anticoagulant. Sex could not be reliably determined because most gonads were undeveloped.

**Fig 1.**
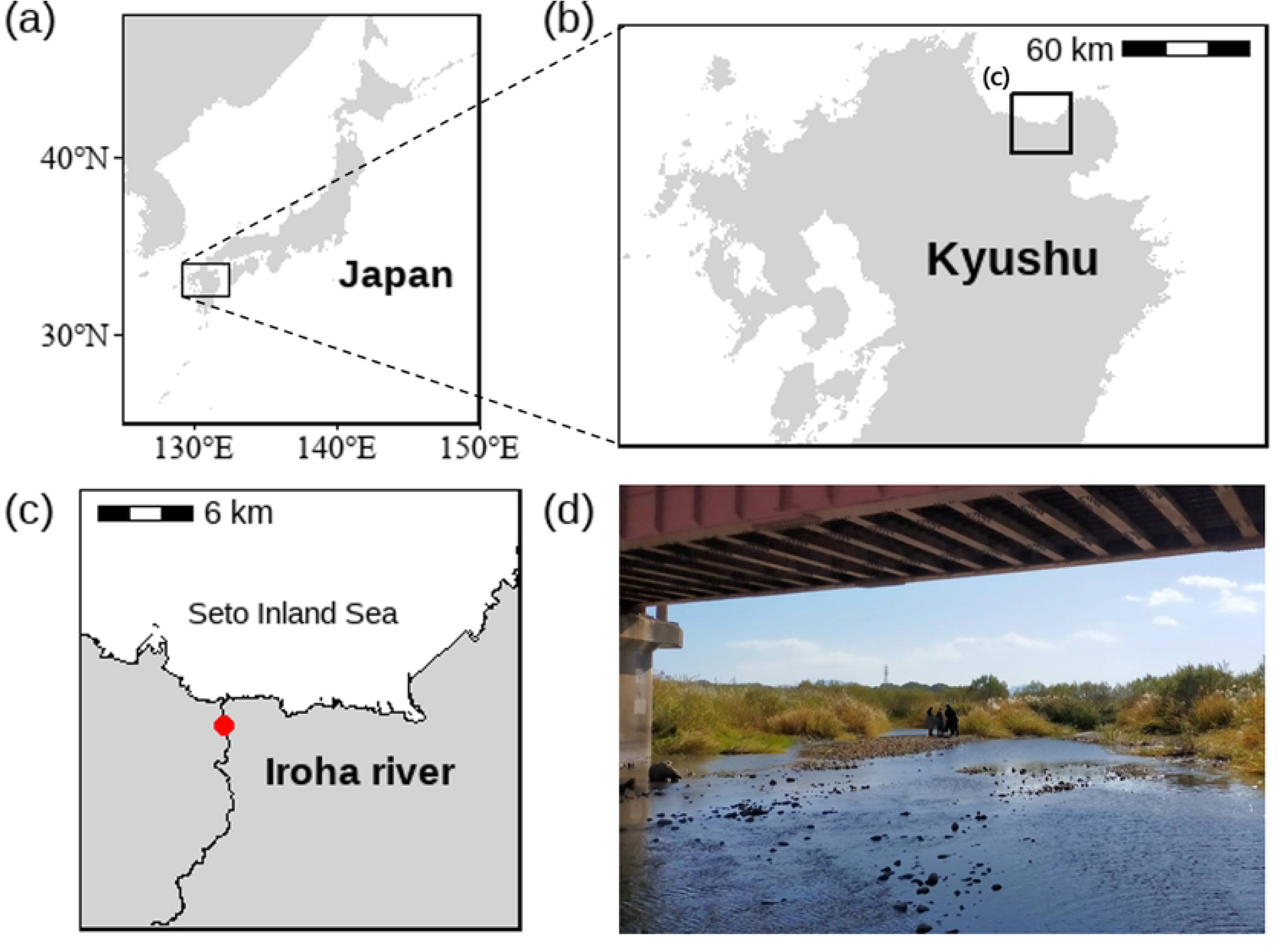
Maps and photograph of the sampling sites. (a) Map of Japan. (b) Map showing Kyushu, Japan. (c) Map of the Iroha River. Japanese eels were collected at the site marked with a red circle (approximately 300 m). (d) Photograph of sampling site in the Iroha River (November 7, 2023).

All procedures were conducted under permits and approvals from Oita Prefecture (Permit No. Kitakyoku toku dai 9 gou) and Kyoto University (Approval Nos. I-202301 and I-202401).

### 2-3. DNA extraction from wild individuals

DNA was extracted using the DNeasy Blood & Tissue Kit (QIAGEN, Venlo, Netherlands) following the manufacturer’s protocol, except that the final elution incubation time at room temperature was extended from 1 to 5 min to increase DNA yield [43]. Sufficient DNA purity (A_260_/A_280_ ratio: approximately 1.6–2.0) and concentration (> 10 ng/µL) were confirmed using a microvolume spectrophotometer (e.g., DS-11, DeNovix Inc., Wilmington, DE, USA). The extracted DNA was stored at −30°C.

### 2-4. Telomere length measurement

TL was measured by qPCR using the Applied Biosystems StepOnePlus Real-Time PCR System (Applied Biosystems, Thermo Fisher Scientific, Waltham, MA, USA). The ribosomal protein L7 gene (*rpl7*; NCBI Gene ID: 118234739) was used as the reference gene, amplified with primers RPL7_F: 5’-GGA AGT TGT TTG CGG CCT TGA A-3’, RPL7_R: 5’-AGG GCA GGG GTC TCT TCA TAC A-3’. For primers targeting telomere sequences, we used tel1b: 5’-CGG TTT GTT TGG GTT TGG GTT TGG GTT TGG GTT TGG GTT-3’ and tel2b: 5’-GGC TTG CCT TAC CCT TAC CCT TAC CCT TAC CCT TAC CCT-3’ [44].

DNA samples were thawed and diluted with molecular biology grade water (Sigma-Aldrich, Merck KGaA, Darmstadt, Germany) to a final concentration of 5 ng/µL prior to telomere length measurement experiments. Each 20 µL qPCR reaction contained 10 µL TB Green® Premix Ex Taq™ II (Tli RNaseH Plus, 2×; Takara Bio Inc., Shiga, Japan), 0.4 µL ROX Reference Dye (50×; Takara Bio), 0.4 µL each of forward and reverse primers (10 µM), 2 µL DNA template (5 ng/µL) or no-template control, and 6.8 µL nuclease-free water. After dispensing, a film (Micro Amp™ Optical Adhesive Film, Thermo Fisher Scientific, Waltham, MA, USA) was placed over the plate for the reaction.

Each sample was run in triplicate, and tissues from the same individual were placed on the same plate. Telomere and reference gene assays were run on separate plates. A five-point standard curve (50, 25, 12.5, 5, 1 ng/µL) was included in duplicate on each plate using liver DNA from Japanese eel as a calibrator to estimate amplification efficiency.

The cycling protocol was 95°C for 30 s, followed by 40 cycles of 95°C for 5 s and 58°C for 30 s, with a final melt-curve step (95°C for 15 s, 60°C for 1 min, and 95°C for 15 s). All samples exhibited a single melt-curve peak, indicating specific amplification. Ct values and amplification efficiencies were calculated using StepOne™ Software v2.3 (Applied Biosystems). Amplification efficiencies were 0.957 ± 0.076 for RPL7 and 1.048 ± 0.116 for telomeric sequences. Relative telomere length (RTL) was calculated according to the Pfaffl’s method [45]:

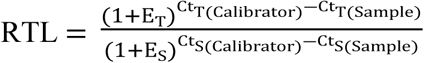

where E_T_ and E_S_ are the amplification efficiencies for the telomere sequence (T) and reference gene (S), respectively, in each plate. Ct_T(Calibrator)_ and Ct_S(Calibrator)_ are mean Ct values for the calibrator, and Ct_T(Sample)_ and Ct_S(Sample)_ are those for each sample.

For quality control, data points outside the standard curve range were excluded. When the SD of Ct values among triplicates exceeded 0.13 [46], the replicate with the largest absolute deviation from the triplicate mean was excluded and the remaining two Ct values were averaged. If SD remained ≥ 0.13 after exclusion, all three values were retained. This quality-control rule was pre-specified and applied symmetrically to both telomeric and reference gene assays. The intra-assay CV was 0.242% for RPL7 and 1.198% for the telomeric sequence; inter-assay CV (*n* = 13 plates) were 2.522% for RPL7 and 5.036% for the telomeric sequence.

### 2-5. Statistical analysis

All statistical analyses were conducted in R v4.3.2 [47] with the statistical significance level at 0.05. RTL data for each tissue did not meet normality (Shapiro–Wilk test, *p* < 0.05). To evaluate the effects of tissue type on RTL and to examine the tissue-specific relationships between RTL and either total length or age, a generalized additive mixed model (GAMM) was constructed using the gamm4 package [48]. The model used a Gamma error distribution with log link function, including tissue type as a fixed effect, smooth terms for total length (or age) interacting with tissue type, and individual ID as a random effect. Two competing models were fitted, one including total length and the other including age as predictors, and compared using Akaike’s information criterion (AIC). Total length was Z-score transformed prior to GAMM fitting to improve model convergence.

To further explore RTL differences among tissues, pairwise Wilcoxon’s signed-rank tests adjusted with Bonferroni correction were performed using the 18 individuals from which all tissues were successfully collected. Additionally, associations in RTL among tissues were assessed using Spearman’s rank correlations with Bonferroni correction.

### 2-6. AI policy statement

Chat GPT (v5.3, Open AI) was used to provide partial support for R code development and troubleshooting, as well as for improving the clarity and wording of the English manuscript.

## Results

### 3-1. FISH analysis

Fluorescence signals for the TTAGGG telomeric sequences were observed at all chromosomal ends of the Japanese eel (2*n* = 38) (Fig 2a). ITSs were detected on three pairs of chromosomes (Fig 2b); however, the fluorescence intensity was weaker than that of the terminal telomeres.

**Fig 2.**
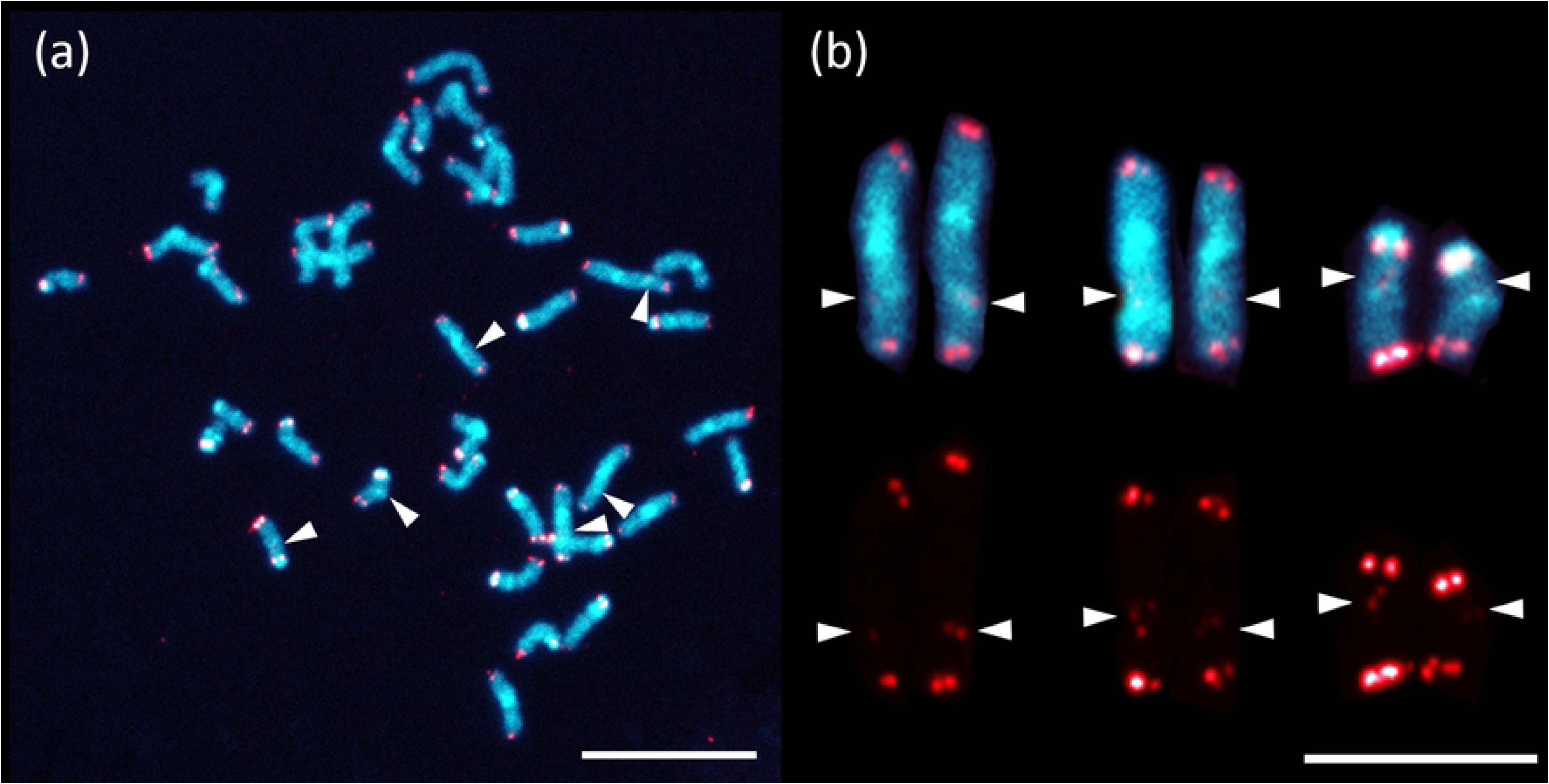
Chromosomal distribution of TTAGGG telomeric sequences (red) of the Japanese eel (*Anguilla japonica*). (a) A metaphase spread. (b) Three chromosome pairs with ITS signals from the metaphase shown in Fig 2a. The upper panel shows chromosomes with detected ITS signals, while the lower panel shows only the telomeric signals (red) extracted from these chromosomes. Arrowheads indicate ITSs signals in both panels. Scale bars: 10 µm in (a), 5 µm in (b).

### 3-2. Tissue differences in RTL

RTL was significantly affected by tissue type (Table 1). Within individuals, caudal fin RTL was generally the longest, exceeding those in all other tissues (*p* < 0.05 for all pairwise comparisons after Bonferroni correction; Fig 3, Fig S1) except blood and liver, where RTL was comparable (*p* = 1.000 for both). Conversely, brain and muscle RTL were significantly shorter than those in all other tissues (*p* < 0.05 for all pair; Fig 3, Fig S1).

**Fig 3.**
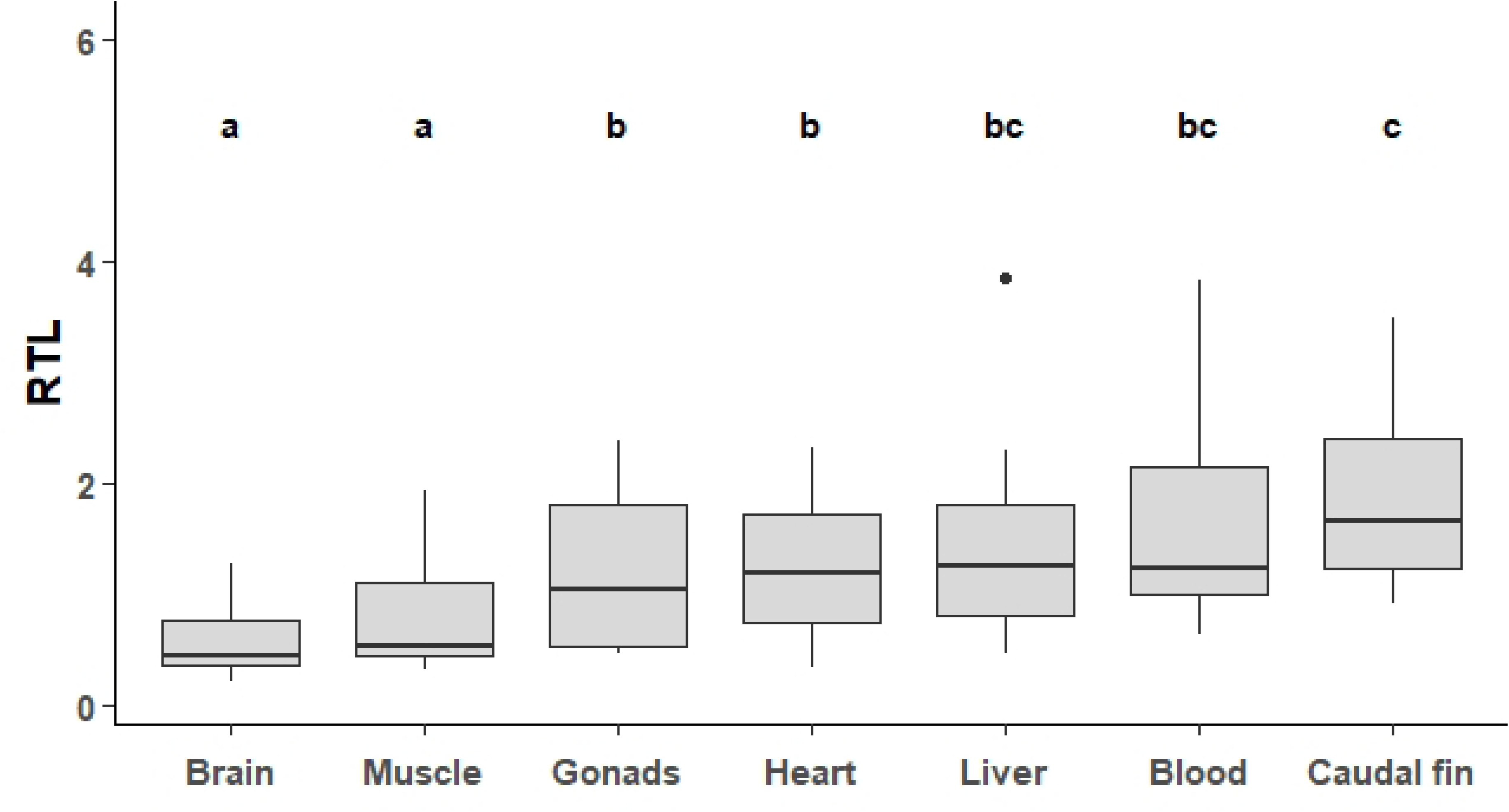
Comparison of relative telomere length (RTL) among tissue types in the Japanese eel (*Anguilla japonica*). Boxplots show the distribution of individuals with complete tissue sets (*N* = 18). Different letters denote significant differences among tissues based on pairwise Wilcoxon’s signed-rank tests with Bonferroni correction.

**Table 1.** Summary of GAMM evaluating tissue effects and tissue-specific smooth terms of total length on RTL (reference tissue: muscle). Bold letter indicates significant.

|  | Estimate | SE | *t* value | *p* value |
| --- | --- | --- | --- | --- |
| **(Intercept)** | -0.232 | 0.079 | -2.928 | **0.003** |
| **Blood** | 0.819 | 0.060 | 13.58 | **< 0.001** |
| **Brain** | -0.125 | 0.059 | -2.113 | 0.035 |
| **Caudal fin** | 0.821 | 0.060 | 13.80 | **< 0.001** |
| **Gonads** | 0.228 | 0.126 | 1.806 | 0.071 |
| **Heart** | 0.297 | 0.060 | 4.979 | **< 0.001** |
| **Liver** | 0.537 | 0.059 | 9.041 | **< 0.001** |
|  | edf | Ref.df | *F* | *p* value |
| **s(Total Length)**<br>**× Muscle** | 2.564 | 2.564 | 5.331 | **0.013** |
| **s(Total Length)**<br>**× Blood** | 2.680 | 2.680 | 6.739 | **0.003** |
| **s(Total Length)**<br>**× Brain** | 2.426 | 2.426 | 10.06 | **< 0.001** |
| **s(Total Length)**<br>**× Caudal fin** | 1.001 | 1.001 | 1.112 | 0.293 |
| **s(Total Length)**<br>**× Gonads** | 1.000 | 1.000 | 0.753 | 0.386 |
| **s(Total Length)**<br>**× Heart** | 1.012 | 1.012 | 0.005 | 0.965 |
| **s(Total Length)**<br>**× Liver** | 1.004 | 1.004 | 2.205 | 0.138 |
| *Random effects* | Variance | SD |  |  |
| **ID** | 0.073 | 0.270 |  |  |
| **Residual** | 0.120 | 0.347 |  |  |

### 3-3. Correlation of RTL among tissues

RTL in muscle significantly positive-correlated with that in brain, heart, liver, blood, and caudal fin (*p* < 0.05 for all; Table 2, Fig 4), but not with that in gonads (*p* = 0.675; Fig S1). Also, RTL in caudal fin significantly positive-correlated with that in all tissues (*p* < 0.05 for all; Table 2, Fig 4) except blood (*p* = 0.266; Fig S2).

**Fig 4.**
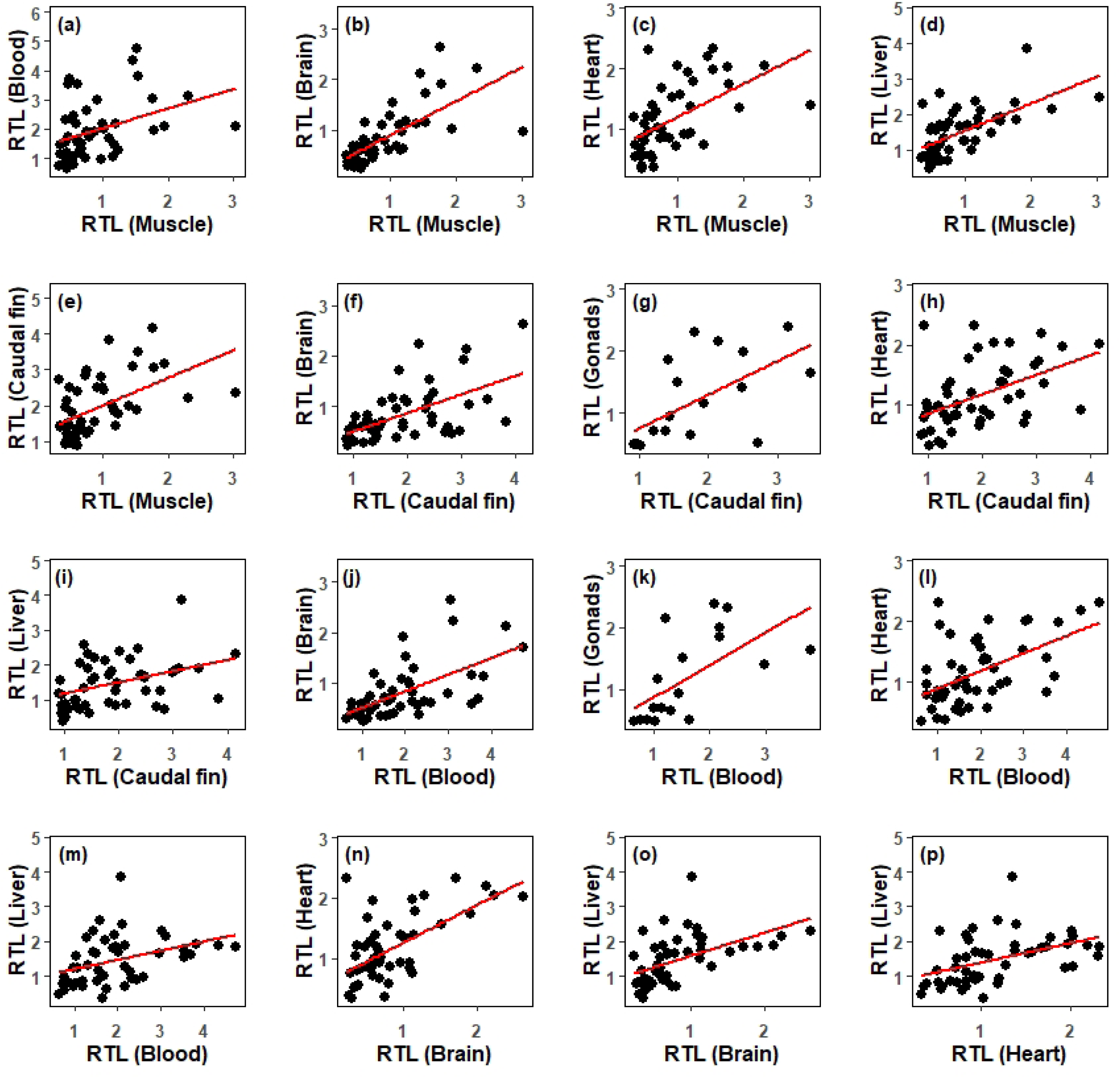
Significant correlations in RTL among tissues (a) Muscle and blood, (b) Muscle and brain, (c) Muscle and heart, (d) Muscle and liver, (e) Muscle and caudal fin, (f) Caudal fin and brain, (g) Caudal fin and gonads, (h) Caudal fin and heart, (i) Caudal fin and liver, (j) Blood and brain, (k) Blood and gonads, (l) Blood and heart, (m) Blood and liver, (n) Brain and heart, (o) Brain and liver, (p) Heart and liver. Correlation coefficients and Bonferroni-adjusted *p*-values are shown in Table 2.

**Table 2.**
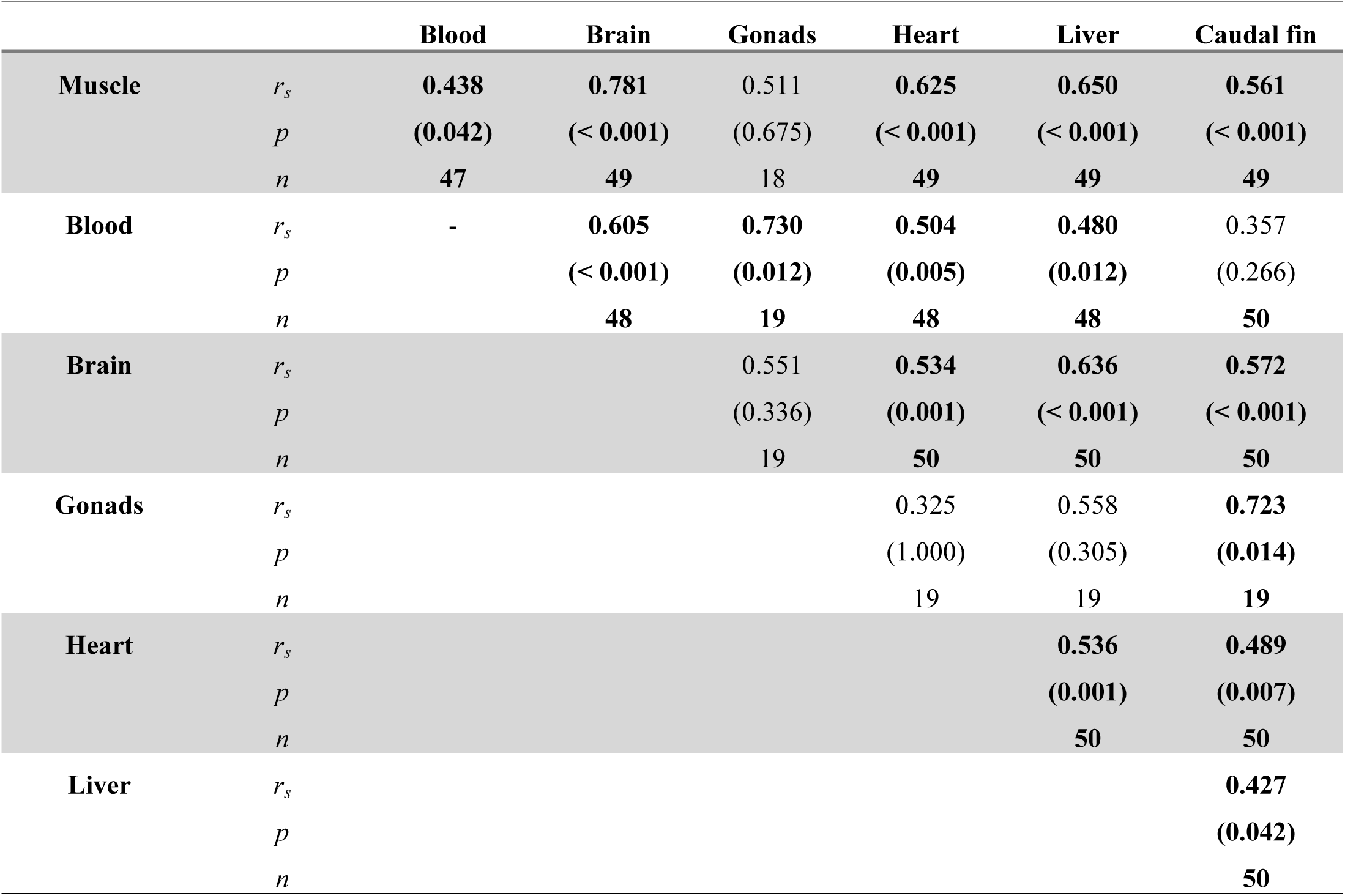
Spearman’s correlation coefficient for RTL among different tissue types and Bonferroni-adjusted p-values. Bold letter indicates significant.

### 3-4. Relationships of RTL with total length and age

Both total length and age were tested as explanatory variables for RTL. The model including total length showed a lower AIC value than the age model (ΔAIC = 18.86). The total length of specimens ranged from 13.9 to 54.1 cm, with an average of 32.1 ± 11.2 cm (SD). A nonlinear relationship between total length and RTL in blood, brain, and muscle tissues (GAMM, *p* < 0.05; Table 1, Fig 5). RTL decreased with increasing total length up to approximately 30 cm in blood and then remained relatively constant, whereas in brain and muscle it exhibited a hump-shaped pattern, peaking at approximately 22 cm and 29 cm, respectively. No significant relationship with total length was detected in the other tissues (*p* > 0.05; Table 1, Fig. S3).

**Fig 5.**
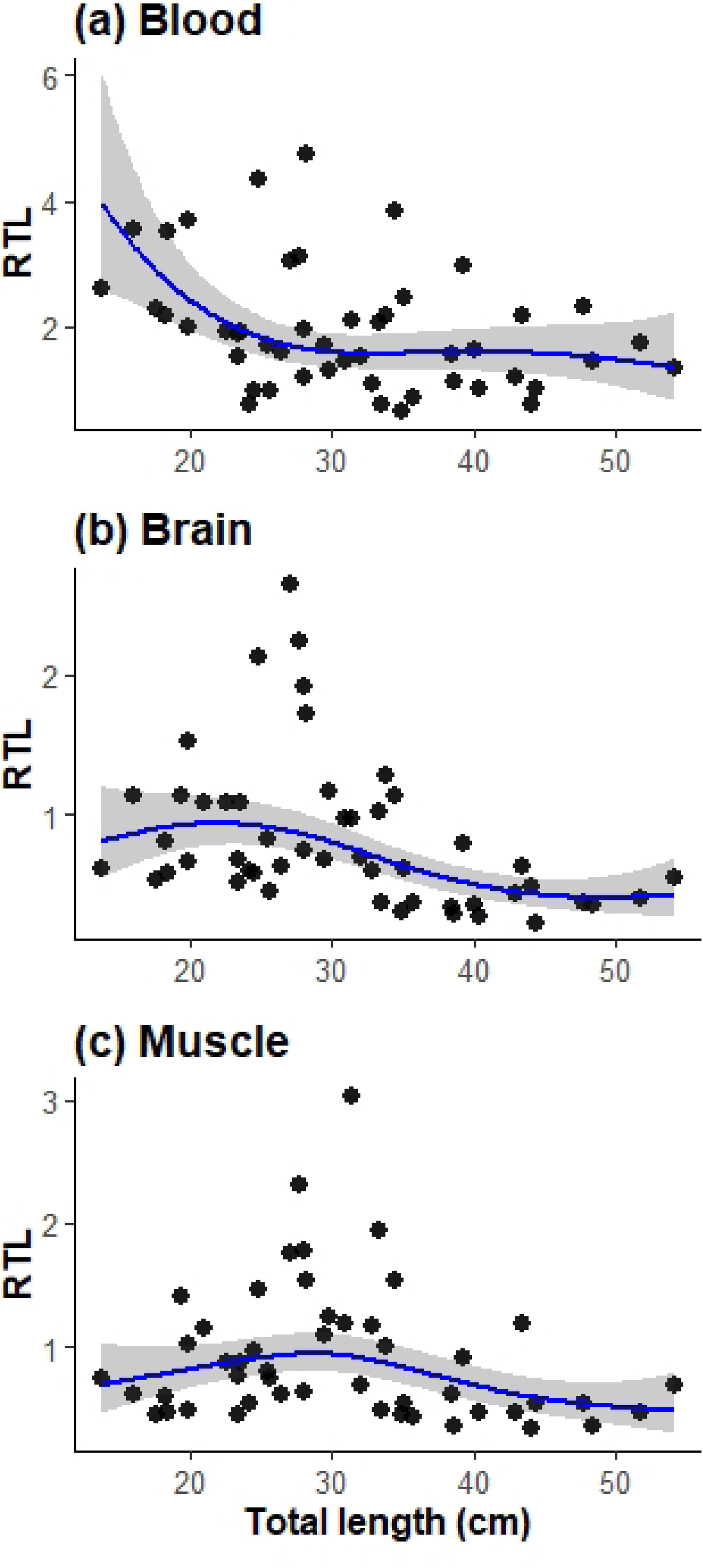
RTL in tissues exhibiting nonlinear effects of total length Black points represent observed values, blue lines indicate fitted curves based on GAMM, and gray shaded areas represent 95% confidence intervals.

In the age-based model, a significant relationship was detected only in brain (*p* < 0.001; Fig. S4), whereas no significant effects were found in other tissues (*p* > 0.05; Table S1, Fig. S4).

## Discussion

The preset study provides the first integrative findings and visualization of chromosomal telomeric sequence distribution, as well as cross-tissue assessments of TL, in Japanese eel. In this study, ITSs were detected on three chromosome pairs by FISH. ITSs have also been reported in the American eel (*A. rostrata*) and the European eel (*A. anguilla*), where telomeric repeats occur not only at chromosome ends but also within the nucleolar organizer region on chromosome 8 [49], suggesting that such ITS-like arrangements may be characteristic of the genus *Anguilla*. In this study, the signal intensity of ITSs was weaker than that of terminal telomeres. Although an excess amount of ITSs may lead to overestimation of TL [40] as qPCR-based TL quantification amplifies ITSs as well as terminal telomeres, our findings suggest that the influence of ITSs on RTL estimates using qPCR is likely limited in this species. Notably, substantial among-tissue variation in RTL was observed. RTL in brain and muscle was relatively short, whereas RTL in blood and caudal fin was markedly long. Tissue-specific TL dynamics has been reported in vertebrates [e.g., 50], and these results show that tissue-specific variation in TL exists in Japanese eel as well. These findings also support the conclusion that the presence of fixed ITSs does not substantially compromise the accuracy of qPCR-based TL measurements.

Cross-tissue comparisons across taxa demonstrate that suitable proxy tissues for TL measurement can be highly species-specific. In humans, dogs, pigs, zebra finches, brown Norway rats (*Rattus norvegicus*), and lizards, blood TL significantly correlated with TL in other tissues [18–23, 51]. Dorsal fin RTL also correlates with other tissues in brown trout [28]. In contrast, some taxa show little or no inter-tissue correlation: for example, wing RTL—not blood RTL—best reflects internal tissues in Egyptian fruit bats [24], and correlations are weak or absent in blind mole-rats and aquaculture cod [30, 51]. These comparisons highlight that the suitable tissue types for TL measurement must be identified on a species-specific basis. Our findings suggest that in Japanese eel, muscle and caudal fin, which are non-lethal and relatively easy to obtain in the field, provide practical, non-lethal proxies for assessing individual telomere dynamics. Importantly, their regenerative capacity enables repeated sampling of the same individuals, which is essential for longitudinal assessments of telomere dynamics at the individual level [39]. This approach, therefore, may facilitate future studies assessing internal physiological costs or responses to external stressors [e.g., 26, 52, 53].

Tissue differences in RTL may arise from variation in cell division, metabolic rates, stem-cell abundance, and telomerase regulation [54–56]. Although tissues with high cell-division rates generally exhibit shorter telomeres [24, 56], Japanese eel showed the opposite pattern: brain and muscle—both largely composed of post-mitotic cells— RTL was short, whereas blood and caudal fin RTL was long. This opposite pattern may reflect tissue-specific telomere regulation in fishes, as regenerative tissues such as fins can maintain telomerase activity [e.g., 57]. In fact, tissue-specific telomerase activity has been confirmed in zebrafish [55, 57].

Alternatively, ecological factors or physiological demands could also contribute to tissue differences in RTL via oxidative stress [4, 5, 58] and tissue-specific metabolic rates [54, 59]. Japanese eels utilize gaps within river substrates as resting habitat [60–62] and are active, top-level predators in river food webs [63]. Sustained whole-body swimming likely elevates muscular oxidative metabolism and ROS exposure [64], whereas spatial and recognition may impose high metabolic demands on the brain [65], both potentially contributing to RTL shortening.

We also found nonlinear relationships between RTL and total length. In blood, RTL declined up to approximately 30 cm in total length, and then stabilized. This pattern is consistent with biphasic telomere loss during early hematopoiesis—rapid shortening during periods of high progenitor turnover, followed by a much slower rate thereafter—reported for human leukocyte lineages and hematopoietic stem-cell systems [66]. As hematopoietic activity is expected to decline with growth, continued but limited telomerase activity in hematopoietic cells may contribute to maintaining RTL at later stages [e.g., 67]. Similar early-life telomere shortening followed by reduced rates of change has also been documented in birds [68], suggesting that such nonlinear dynamics may represent a common feature of telomere regulation during early growth. Nevertheless, comparable evidence in fish remains limited, and our findings may provide novel insight into telomere dynamics in aquatic ectotherms.

In muscle and brain, hump-shaped nonlinear effects were observed where RTL peaked at 20–30 cm in total length. In Japanese eel, sex differentiation begins at approximately 30 cm in total length [69]. Since sex steroids can upregulate telomerase activity via TERT regulation [70–72], the relatively longer RTL observed in muscle and brain around 25 cm in total length may be consistent with hormonally mediated increases in telomerase accompanying the onset of sex differentiation. Moreover, skeletal muscle and the central nervous system are generally well-recognized sex steroid-responsive tissues [e.g., 73, 74], which could explain why longer RTL was evident in these tissues but not others.

Several limitations of this study should be acknowledged. First, although we interpret tissue-specific RTL patterns in light of ecological and physiological hypotheses, direct measurements of telomerase activity, oxidative stress, and metabolic rates across tissues in Japanese eel are currently lacking. Second, although we analyzed multiple tissues from wild individuals, sample sizes for both the smallest and largest size classes were limited, which reduces the precision of estimated nonlinear RTL–growth relationships. Notably, the smallest individual from which gonads could be collected measured only 29.8 cm in total length, and sampling around the onset of sex differentiation (approximately 30 cm in total length; [69]) was not sufficient. Additionally, sex determination was impossible for most individuals. Therefore, it is difficult to fully evaluate growth-related effects on gonads, tissues most directly associated with sexual differentiation, or to assess the potential influence of sex or sex steroids on tissue-specific telomere dynamics. Addressing these limitations will improve our understanding of how intrinsic and extrinsic factors shape telomere dynamics in this species.

In conclusion, this study represents the first application of FISH in Japanese eel, confirming that ITSs are limited in this species. Furthermore, our study is the first example to measure TL from multiple tissues in wild eels. This study revealed that TLs in muscle and caudal fin can represent the individual telomere dynamics, allowing future studies to track temporal changes in physiological condition and to evaluate telomere responses to environmental variation or stress. In recent years, telomere research has become increasingly important in non-model species with diverse life histories [10, 24]. Future studies are needed to examine telomere dynamics and inter-tissue relationships across diverse taxa to support the use TL as an indicator of physiological and conservation status.

## Acknowledgements

This study was supported by JST FOREST (Grant Number JPMJFR2171), the River Fund (Grant Numbers 2024-5211-032 and 2025-5211-046), and JSPS KAKENHI (Grant Number 24K08950). We thank the Medical Research Support Center, Graduate School of Medicine, Kyoto University, for providing access to experimental equipment. We are grateful to Mr. Kanato Ogiso (Graduate School of Agriculture, Kyoto University), Dr. Mayu I. Ogawa (currently at JAMSTEC, formerly Graduate School of Agriculture, Kyoto University) and Ms. Haruka Nakajin (Graduate School of Agriculture, Kyoto University) for their generous assistance in sampling. We also thank Dr. Yuichi Mizutani (Lecturer, Faculty of Environmental Science, University of Human Environments) for his valuable advice on data analysis. Finally, we express our sincere appreciation to all members of the Fisheries and Environmental Oceanography Laboratory, Graduate School of Agriculture, Kyoto University, for their continuous support throughout this study.

## Author Contributions

Conceptualization: Yuta Moriguchi and Satoko S. Kimura.

Data curation: Yuta Moriguchi and Yoshinobu Uno.

Formal analysis: Yuta Moriguchi. Funding acquisition: Satoko S. Kimura.

Investigation: Yuta Moriguchi, Satoko S. Kimura, Manabu Kume, Junichi Takagi, and Yoshinobu Uno.

Methodology: Yuta Moriguchi, Satoko S. Kimura, Manabu Kume, Junichi Takagi, Yoshinobu Uno, Junko Kanoh, and Hiromichi Mitamura.

Project administration: Satoko S. Kimura and Hiromichi Mitamura.

Validation: Yuta Moriguchi and Satoko S. Kimura.

Visualization: Yuta Moriguchi.

Writing – original draft: Yuta Moriguchi and Yoshinobu Uno.

Writing – review & editing: Yuta Moriguchi, Satoko S. Kimura, Manabu Kume, Junichi Takagi, Yoshinobu Uno, Junko Kanoh, and Hiromichi Mitamura.

## Supporting information

**Fig S1.**
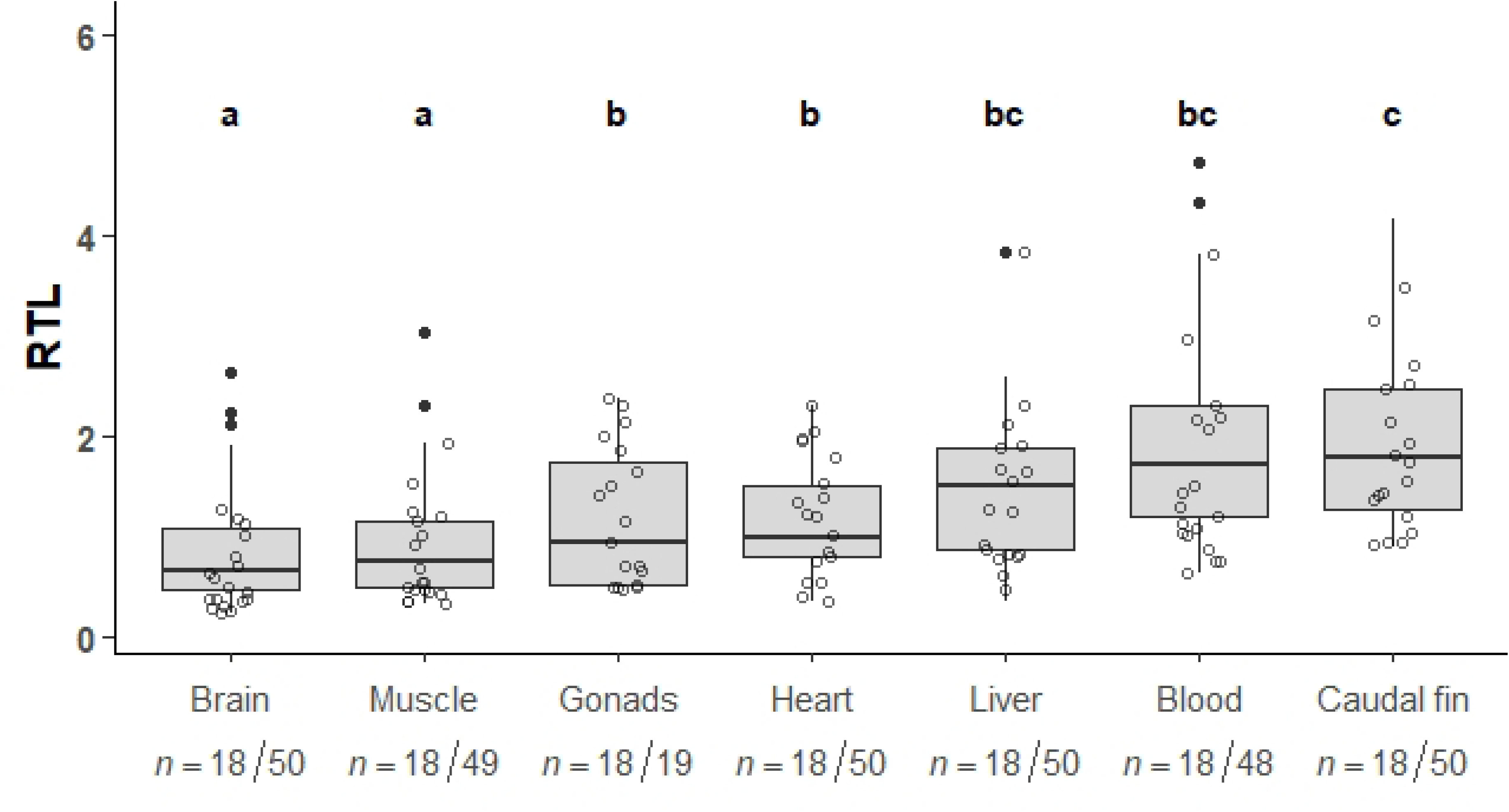
Comparison of relative telomere length (RTL) among tissue types in the Japanese eel (*Anguilla japonica*). Boxplots show the distribution of all available samples, while white circles indicate individuals with complete tissue sets used for paired comparisons. The numbers below each tissue indicate the number of paired samples / total samples. Different letters denote significant differences among tissues using pairwise Wilcoxon’s signed-rank tests with Bonferroni-correction.

**Fig S2.**
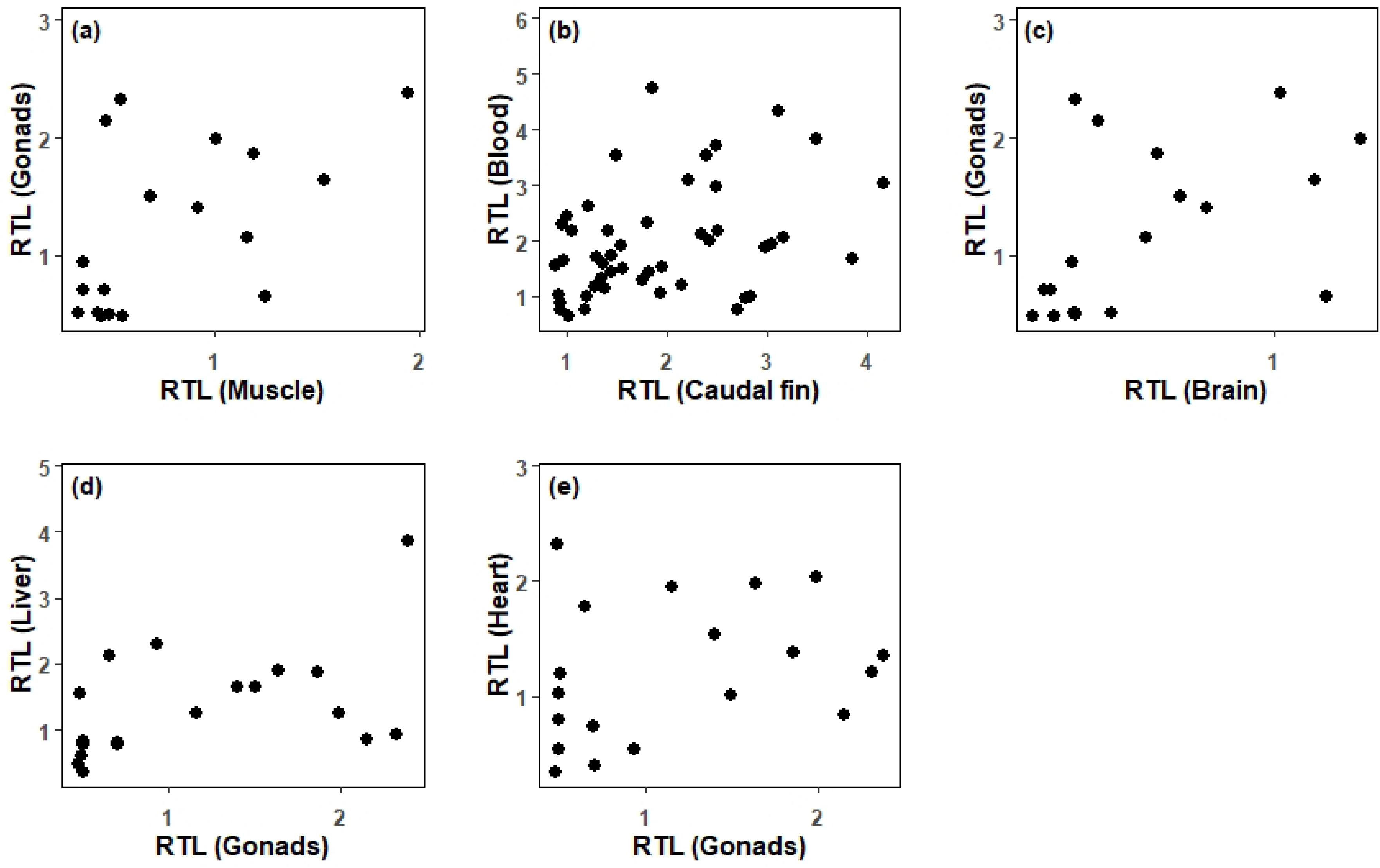
The relationship between RTLs in tissues where no significant correlation was observed. (a) Muscle-Gonads, (b) Caudal fin-blood, (c) Brain-Gonads, (d) Gonads-liver, (e) Gonads-heart.

**Fig S3.**
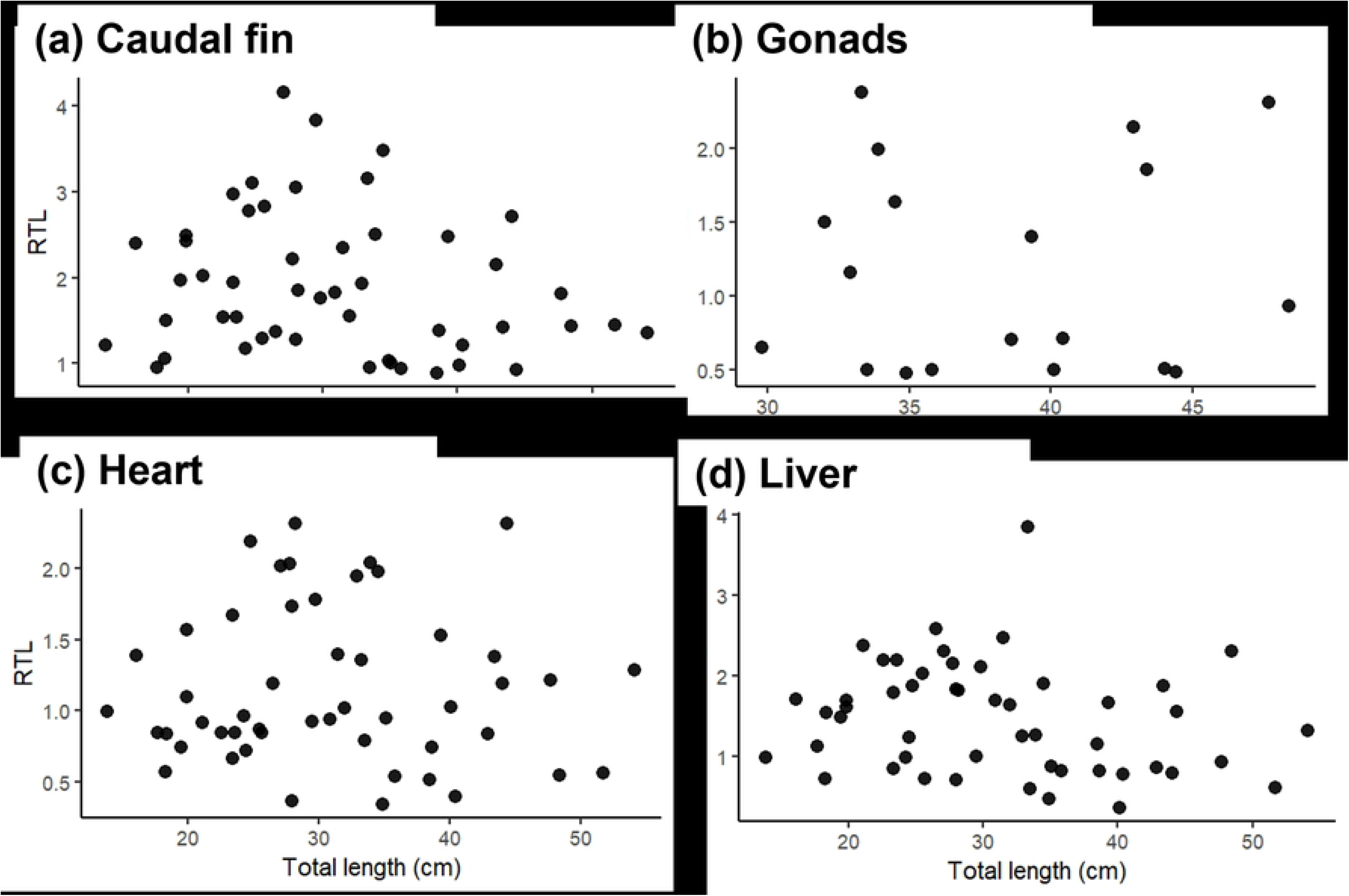
The relationship between total length and RTL of each tissue except blood, brain, and muscle.

**Fig S4.**
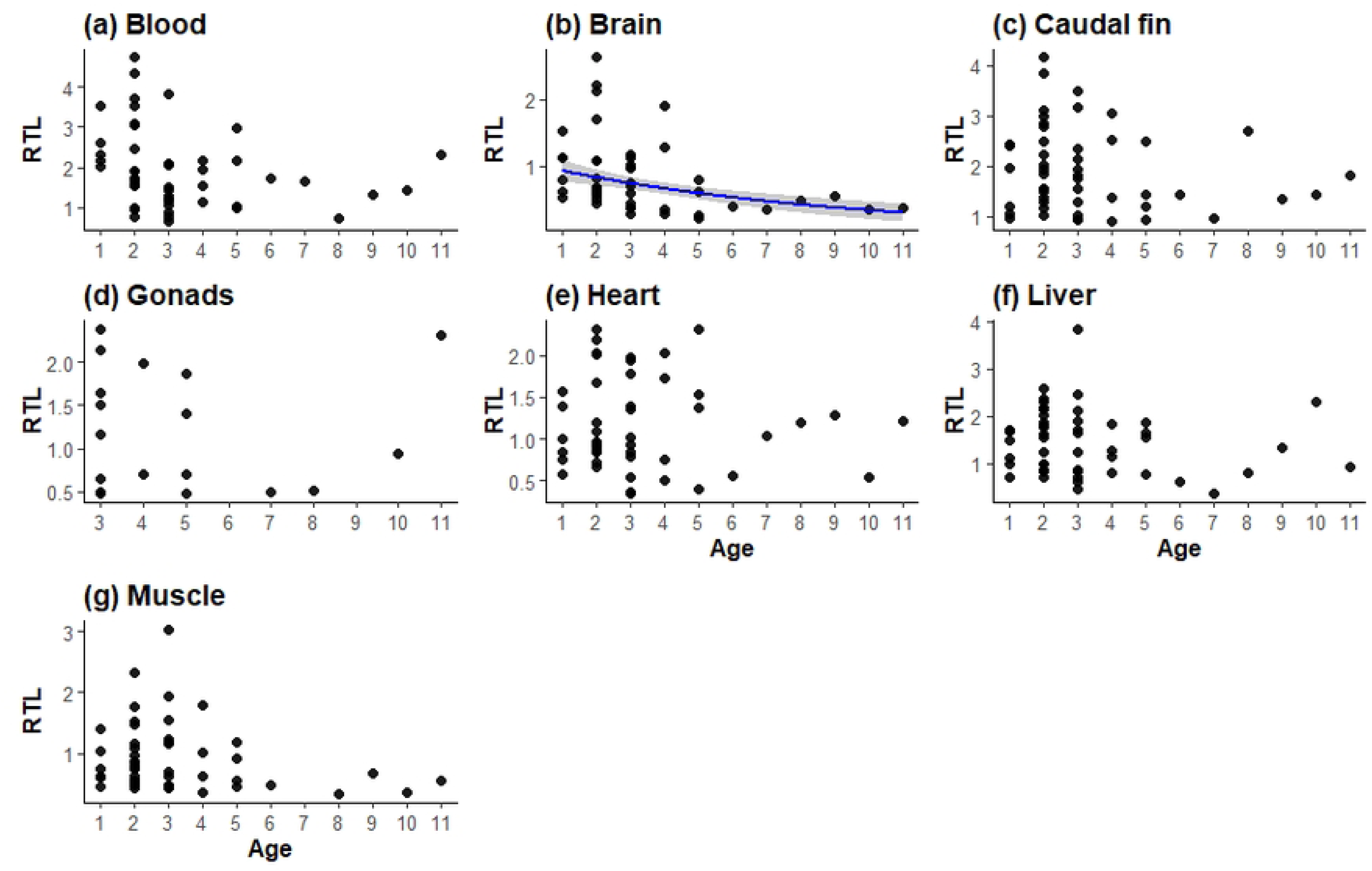
The relationship between age and RTL of each tissue. Black points represent observed values, blue lines indicate fitted curves based on GAMM (age-based model), and gray shaded areas represent 95% confidence intervals.

**Table S1.** Summary of the model-based evaluation of the association between RTL and age (reference tissue: muscle). Bold letter indicates significant.

|  | Estimate | SE | *t* value | *p* value |
| --- | --- | --- | --- | --- |
| **(Intercept)** | -0.223 | 0.080 | -2.768 | **0.006** |
| **Blood** | 0.823 | 0.063 | 13.09 | **< 0.001** |
| **Brain** | -0.115 | 0.062 | -1.868 | 0.062 |
| **Caudal fin** | 0.816 | 0.062 | 13.20 | **< 0.001** |
| **Gonads** | 0.302 | 0.193 | 1.565 | 0.118 |
| **Heart** | 0.291 | 0.062 | 4.698 | **< 0.001** |
| **Liver** | 0.533 | 0.062 | 8.624 | **< 0.001** |
|  | **edf** | **Ref.df** | **F** | **p value** |
| **s(Total Length)**<br>**× Muscle** | 1.577 | 1.577 | 4.474 | 0.055 |
| **s(Total Length)**<br>**× Blood** | 1.795 | 1.795 | 2.391 | 0.055 |
| **s(Total Length)**<br>**× Brain** | 1.000 | 1.000 | 14.85 | **< 0.001** |
| **s(Total Length)**<br>**× Caudal fin** | 1.000 | 1.000 | 0.527 | 0.468 |
| **s(Total Length)**<br>**× Gonads** | 2.010 | 2.010 | 0.102 | 0.791 |
| **s(Total Length)**<br>**× Heart** | 1.028 | 1.028 | 0.009 | 0.951 |
| **s(Total Length)**<br>**× Liver** | 1.000 | 1.000 | 0.797 | 0.373 |
| **Random effects** | **Variance** | **SD** |  |  |
| **ID** | 0.076 | 0.276 |  |  |
| **Residual** | 0.130 | 0.360 |  |  |

